# Reconstructing the Huang Surname and Its Related Lineages: A Comprehensive Analysis of Molecular Genetics and Historical Genealogies

**DOI:** 10.1101/2025.02.17.638729

**Authors:** Shi Huang

## Abstract

The origin of the Huang surname is complex, but according to historical texts such as *Shiji*, it can be traced back to the Yellow Emperor (Huang Di) and his descendant, Bo Yi, who played a key role in controlling the floods during the Xia Dynasty and was granted the surname Ying. Bo Yi had two sons who each established a state, one named Huang and the other Xu. After the fall of Huang, its people adopted Huang as their surname and migrated to Hubei. Similarly, after the fall of Xu, its people adopted Xu as their surname, spreading throughout Jiangsu. Further descendants of Bo Yi also gave rise to 12 other surnames, including the Liang surname. Given that the narratives of ancient texts are often regarded as legendary, we aim to independently verify the authenticity of the historical records by analyzing the paternal haplotypes of the Huang surname and its related surnames using scientific methods. We analyzed the surname genetic data of 207,333 Han Chinese males published by 23Mofang and combined it with independent sequencing research. In 189 Y chromosome haplotype branches, each containing at least five individuals with the surnames Huang, Xu, or Liang, and formed around 4,000 years ago, we analyzed the proportion of samples with the Huang surname from Hubei among all samples nationwide with the Huang surname. Branches with a high proportion in Hubei are likely to represent the Hubei-specific lineage of the Huang surname. We similarly analyzed the proportion of samples with the Xu surname in the Jiangsu area and of samples with the Liang surname in the Hebei area. We further applied two criteria to identify the ancestral genotype of a specific surname in a given region: first, the surname should appear at a high frequency among all surnames within that genotype; second, the proportion of samples of the region among nationwide samples should be high. The results showed that the Hubei-specific Huang surname likely carries the MF14296/MF15137 haplotype downstream of the supergrandfather O-M117-F8-F2137-A16635-A16636. Our independent sampling study found that A16636 is the dominant haplotype among the Huang surname population in nine neighboring Huang surname villages around Luohansi in Huangpi, Hubei. The Jiangsu-specific Xu surname likely carries the MF38096/SK1726 haplotype under the A16636 sister branch F15823, while the Hebei-specific Liang surname likely carries the F15823 related MF14287/MF15398/MF14963 haplotype. This conclusion aligns with the historical records, providing strong genetic evidence that supports the authenticity of these ancestral claims regarding the Huang and Xu states and the Xia dynasty.

## Introduction

Chinese Han surnames began to form around 5,000 years ago (1–3). By 4,000 years ago, the eight major ancient surnames existed, and they are considered the ancestors of today’s approximately 23,000 surnames (4). Among them, the Ying surname is believed to be the ancestor of 14 present day surnames, including Huang, Xu, Liang, Zhao, Qin, Jiang, Ge, Zhai, Gu, Ma, Xiao, Zhong, Miao, and Fei. Five of these surnames are among the top 22 most common surnames in China today, including Huang (ranked 7th), Zhao (8th), Xu (11th), Ma (13th), and Liang (22nd).

The great historian Sima Qian of the Han dynasty, living from around 145 BC to 86 BC and arguably the greatest historian in Chinese history, in his monumental work *Records of the Grand Historian* (*Shiji*), provides the following account regarding the origins of the Qin state: "The ancestors of the Qin were of the Ying surname. Later, they were granted fiefs, adopting the names of the states as their surnames, including the Xu, Tan, Ju, Zhong Li, Yun Yan, Tuqiu, Jiang Liang, Huang, Jiang, Xiu Yu, Bai Ming, Fei Lian, and Qin. Among these, the Qin, whose ancestor Zao Fu was granted the fief of Zhao Cheng, also adopted the Zhao surname." This shows that, among the 14 ancient sub-surnames of the Ying clan recorded in *Shiji*, six surnames, including Xu, Jiang Liang (Liang), Huang, Qin, and Zhao, have clear corresponding surnames today.

According to historical texts such as the *Ancient Bamboo Annals* (*Guben Zhushu Jinian*), the Huang surname originated in the eastern part of Inner Mongolia, south of the Yanshan Mountains, in the region of the Liao River’s western source—now the Xar Moron (Xilamulun) River (formerly known as Huang Shui). This area was home to a tribe called the Huang Yi, who were descendants of Shao Hao, the eldest son of the Yellow Emperor (Huang Di). The Huang Yi tribe later migrated southeastward, passing through the Huang Shan and Huang Qiu areas of Hebei and eventually entering the Shandong Peninsula, where they joined the Eastern Yi group (Dong Yi), becoming one of the Nine Yi tribes.

According to *Shiji*, during the reign of Emperor Shun, Bo Yi, the leader of the Eastern Yi tribe and a descendant of Shao Hao, was granted the surname Ying by Emperor Shun for his role in assisting King Yu of the Xia Dynasty in controlling the floods. Bo Yi had two sons, and in the early 21st century BCE (also said to be around 2148 BCE), his eldest son, Da Lian, established the state of Huang in Longgu Township, 12 miles west of Ding Cheng in Guang Zhou (present-day Huang Chuan, Henan). The ruling family of Huang lasted until 648 BCE, when the state was destroyed by the Chu state. After the fall of Huang, the people of the state adopted the name of their homeland as their surname.

As for the reason why the state was named Huang, it is likely related to the origin of Bo Yi or Shao Hao in the ancient Huang Yi tribe. This tribe liked to use "Huang" to name the places where they lived. For example, in Shandong, Henan, Hubei, and other regions, there are many place names with "Huang" in them, such as Huang County in Shandong, Huangchuan County in Henan, and Huangpi District in Hubei. These place names are probably connected to the migration and settlement history of the Huang Yi tribe. Additionally, it may be related to the fact that Da Lian was the eldest son. The eldest son often carries the family traditions and expectations, a tradition that may have originated very early in Chinese history.

Bo Yi’s second son, Ruo Mu, established the state of Xu (present-day Suqian, Jiangsu). The state continued under this hereditary line until around 500 BCE, when it was destroyed by the Wu state. The passage from *Tongzhi·Shizu Lue* by Zheng Qiao of the Southern Song Dynasty about the Ruo Mu clan reads as follows: "Bo Yi assisted Yu in controlling the floods and achieved great merit, for which his son Ruo Mu was granted land in Xu. His descendants took the name of the state as their surname."

Among the fourteen branches of the Ying surname, only the Huang and Xu surnames directly descend from the sons of Boyi. The other surnames come from more distant descendants of Bo Yi. For instance, Bo Yi’s 14th-generation descendant, Zao Fu, is recorded as the ancestor of the surname Zhao and the surname Ma.

Additionally, according to the historical records in the *Shiji*, *Tongzhi·Shizu Lue,* and other texts, one of Bo Yi’s descendants, known as Fei Zi (who passed away in 858 BCE), was tasked by King Xian of the Zhou Dynasty with overseeing the care of horses and was granted the fief of Qin, becoming known as Qin Ying. Fei Zi’s great-grandson, Qin Zhong, had a younger son who was enfeoffed in Liang, where he established the state of Liang (located in present-day Hancheng, Shaanxi) and was titled Liang Kangbo. Later, the Qin state destroyed the state of Liang.

After the state of Huang was destroyed by the Chu state, the Huang surname group spread, primarily to nearby Hubei. In the north eastern part of Hubei, the cities of Huangpi, Huangan (now Hongan), Huanggang, Huangshi, and Huangmei are arranged from west to east. Among them, Huangpi and Huanggang have a close relationship with the state of Huang(5). Huangpi was once part of the territory controlled by the state of Huang and is located relatively close to the ruins of Huang State in Huangchuan, Henan. Huanggang takes its name from Huanggang Mountain, and according to the *Explanation of County Names* (*Jin Xian Shi Ming*), "To the northwest is Huanggang Mountain, which is named after the ancient state of Huang." Therefore, exploring the origin and development of the Huang surname in Hubei is of great significance for understanding the origins of the Huang surname, the history of the Huang state, and even the history of the Xia Dynasty.

The passage from *Tongzhi·Shizu Lue* reads: "From Ruo Mu to King Yan of Xu, there were 32 generations, until the state was destroyed by the Zhou. The son of the royal family, Zong, was later granted the title of Xu. In the 30th year of Emperor Zhao’s reign, the state was destroyed by the Wu. The descendants took the name of the state as their surname." This passage describes the history of the Xu state, which passed through 32 generations from Ruo Mu in the Xia Dynasty to King Yan of Xu during the Zhou Dynasty. After the Xu state was destroyed by the Wu, the people of the state adopted "Xu" as their surname. The Xu surname group largely remained in Jiangsu after the fall of the Xu state, with the Jiangsu-Shanghai area being the region with the highest frequency of Xu surnames today. Therefore, exploring the origin and development of the Xu surname in the Jiangsu-Shanghai area is of great significance for understanding the origin of the Xu surname, the history of the Xu state, and even the history of the Xia Dynasty. Additionally, it can provide strong corroborative evidence for the origin of the Huang surname as well.

After the fall of the state of Liang, most of its descendants fled to the Jin state, adopting the name of their former country as their surname. The Jin state at the time included the regions of Shanxi and Hebei. Therefore, exploring the history of the Liang surname in the Shanxi and Hebei areas is of great significance for understanding the origin of the Liang surname and the history of the state of Liang. It can also provide supplementary evidence for the origins of the Huang and Xu surnames.

The records in historical texts such as *Shiji* about the Huang surname, the state of Huang, the Xu surname, the state of Xu, Bo Yi, and the Xia Dynasty, if accurate, should theoretically be verifiable through independent research in different fields. Molecular genetics, as a scientific method, can trace the ancestors of surnames and estimate their formation periods, thus providing a possibility for independently verifying the historical accounts of surname origins. The primary aim of this study is to use surname genetics to demonstrate the origins of the Huang surname and related surnames, and to explore the consistency between the molecular genetics findings and the historical records in texts like the *Shiji*.

## Materials and Methods

### Analysis of 23Mofang data

The genetic testing company 23Mofang in Chengdu China has publicly released paternal haplotype data from 220,000 individuals (as of November 8, 2024), including 207,333 Han Chinese individuals, some of which include high-throughput sequencing data. We analyzed these data using the filtering function on the 23Mofang website, which allows for sorting by factors such as surname and region.

For each major paternal haplotype branch of the Huang, Xu, or Liang surnames with at least five samples, dating back around 4,000 years, we analyzed the proportion of Huang surname samples from Hubei among nationwide Huang samples within the same branch. A higher proportion in a branch would suggest it could be the most common or representative Huang surname branch in Hubei, and therefore, likely the one that originally came from the State of Huang and migrated to Hubei. This branch would be the most consistent with historical records about the origin of the Huang surname.

Similarly, we analyzed the proportion of Xu surname samples from Jiangsu and Shanghai (Su-Hu) among nationwide Xu samples within the same branch. A higher proportion here would indicate that this branch is the most representative of the Xu surname in the Su-Hu region.

Additionally, we analyzed the proportion of Liang surname individuals from Shanxi or Hebei among the nationwide Liang samples within the same branch. A higher proportion in this case would indicate that this branch is the most representative of the Liang surname in Shanxi or Hebei.

This approach, based on molecular genetics, independently and impartially explores the origins of the Huang, Xu, and Liang surnames. If the results do not match the historical records, it would challenge the accuracy of those records. If the results align, it would support the reliability and authenticity of the historical accounts.

For the branches identified above, we further analyzed whether the surnames Huang, Xu, or Liang in these branches have a higher proportion compared to all other surnames in the same branch nationwide. Only those with a higher proportion are considered the most likely ancestral branches of the given surname. Additionally, we analyzed the proportion of samples from the respective regions compared to the national total, and branches with a higher proportion are regarded as the most likely region-specific branches. Only those branches that satisfy all three of the above conditions are considered the most regionally representative and most likely to include the ancestral lineage of the surname in that region.

As a comparison for identifying the Hubei-specific Huang surname branch, we also analyzed whether there are similar Jiangsu-specific Huang surname branches.

If there are two or more samples with the same surname, from the same district, and associated with the same account, they are considered to be related, and only one individual is counted. Additionally, key data has been verified with relevant staff members at the 23Mofang company.

Moreover, since most of the 23Mofang clients do not have high-throughput sequencing or lack downstream information for A16636, we sponsored high-throughput sequencing for seven related clients, including three of the six Huang surname individuals who are positive for A16636.

### Sample Collection, DNA Extraction, and Sequencing

We collected saliva samples from seven individuals with the Huang surname, all of whom have ancestral roots in Huangpi, Wuhan, from three different villages. The saliva DNA extraction kit was purchased from Norgen Biotek. DNA extraction, PCR amplification, and PCR fragment sequencing were conducted following standard protocols. The sequencing targeted the A16636 variant, located at position 8,187,115 on the Y chromosome of the Genome Reference Consortium Human Build 37 (GRCh37). PCR forward primer: 5’-TGACACATTGCTGAGCCCAA-3’, reverse primer: 5’-TGTGAGCCAAAACACCTACCA-3’.

Additionally, we collected a blood sample from another Huang surname individual with ancestral roots in Huangpi, Wuhan, and performed whole-genome deep sequencing, which was completed by Novogene. The Y-chromosome sequence data was uploaded to the 23Mofang company, where it was integrated into their database.

All participants provided informed consent.

## Results

### Identification of the Hubei-specific Paternal Branch of the Huang Surname

The genetic sequencing company 23Mofang has released paternal haplotype data for 207,333 Han Chinese males (data as of November 18, 2024), among which Huang surnamed individuals account for 0.024 (i.e., 4,957 individuals). This proportion closely matches the 0.022 recorded in the 2010 national census for the Huang surname population in China (Table 1). Additionally, the distribution of Huang surname individuals in Guangdong and Hubei provinces in the 23Mofang data also aligns with the census results. Specifically, the proportion of Huang surnamed individuals in Guangdong in the data is 0.234, which is similar to the 0.195 found in the census, and the proportion in Hubei is 0.047, which is close to the 0.059 recorded in the census. Therefore, the distribution of the Huang surname in the 23Mofang dataset generally aligns with the census data, without any notable overrepresentation or underrepresentation of Huang surnamed individuals. As a comparison, the national proportion of individuals with the Chen surname in the dataset is 0.05, which corresponds to the 0.0453 found in the national census. In Hubei, the proportion of Chen surnamed individuals is 0.039 in the dataset, which is close to the census proportion of 0.0514, while in Guangdong, the proportion is 0.154 in the dataset, which aligns with the 0.174 found in the census.

**Table 1.** Distribution of Huang and other surnames in Hubei and Guangdong for certain haplotypes in 23Mofang data dated Nov 18, 2024.

| Haplotype | Year BP | Surname | # Han nationwide |  | Proportion | # samples of a surname |  | Ratio of provincial v nationwide |  |
| --- | --- | --- | --- | --- | --- | --- | --- | --- | --- |
|  |  |  | One surname | All surnames |  | Hubei | Guangdong | Hubei | Guangdong |
| A0-T | 149960 | Huang | 4957 | 207333 | 0.024 | 234 | 1159 | 0.047 | 0.234 |
|  |  | Chen | 11529 |  | 0.056 | 446 | 1776 | 0.039 | 0.154 |
| M117 | 13870 | Huang | 993 | 39381 | 0.025 | 40 | 258 | 0.040 | 0.260 |
|  |  | Chen | 2361 |  | 0.060 | 73 | 439 | 0.031 | 0.186 |
| F8 | 7250 | Huang | 934 | 37064 | 0.025 | 36 | 261 | 0.039 | 0.279 |
|  |  | Chen | 2267 |  | 0.061 | 70 | 427 | 0.031 | 0.188 |
| ACT2839 | 7250 | Huang | 934 | 35923 | 0.026 | 35 | 250 | 0.037 | 0.268 |
| F438 | 6800 | Huang | 184 | 12690 | 0.014 | 13 | 29 | 0.071 | 0.158 |
| F14366 | 6790 | Huang | 166 | 11857 | 0.014 | 13 | 27 | 0.078 | 0.163 |
| Y17728 | 6610 | Huang | 159 | 11357 | 0.014 | 13 | 24 | 0.082 | 0.151 |
| ACT7481 | 6590 | Huang | 60 | 5052 | 0.012 | 7 | 14 | 0.117 | 0.233 |
| F1754 | 6530 | Huang | 57 | 4927 | 0.012 | 7 | 14 | 0.123 | 0.246 |
| F26655 | 6510 | Huang | 2 | 295 | 0.007 | 0 | 0 | 0.000 | 0.000 |
| F2643 | 5630 | Huang | 52 | 4509 | 0.012 | 7 | 14 | 0.135 | 0.269 |
| F2137 | 5350 | Huang | 52 | 4509 | 0.012 | 7 | 14 | 0.135 | 0.269 |
| A16635 | 5330 | Huang | 44 | 3110 | 0.014 | 7 | 11 | 0.159 | 0.250 |
|  |  | Chen | 113 |  | 0.036 | 3 | 24 | 0.027 | 0.212 |
|  |  | Wang | 233 |  | 0.075 | 7 | 10 | 0.030 | 0.043 |
|  |  | Zhang | 251 |  | 0.081 | 7 | 19 | 0.028 | 0.076 |
|  |  | Li | 198 |  | 0.064 | 11 | 16 | 0.056 | 0.081 |
|  |  | Liu | 319 |  | 0.103 | 9 | 64 | 0.028 | 0.201 |
|  |  | Zhou | 81 |  | 0.026 | 1 | 18 | 0.012 | 0.222 |
|  |  | Yang | 74 |  | 0.024 | 2 | 4 | 0.027 | 0.054 |
|  |  | Xu | 38 |  | 0.012 | 0 | 5 | 0.000 | 0.132 |
|  |  | Wu | 49 |  | 0.016 | 3 | 3 | 0.061 | 0.061 |
|  |  | Zhao | 76 |  | 0.024 | 2 | 1 | 0.026 | 0.013 |
|  |  | Zhu | 43 |  | 0.014 | 2 | 6 | 0.047 | 0.140 |
| A16636 | 4320 | Huang | 6 | 735 | 0.008 | 3 | 0 | 0.500 | 0.000 |
|  |  | Chen | 17 |  | 0.023 | 0 | 1 | 0.000 | 0.059 |
|  |  | Li | 69 |  | 0.094 | 3 | 1 | 0.043 | 0.014 |
| M F14296 | 3580 | Huang | 6 | 56 | 0.107 | 3 | 0 | 0.500 | 0.000 |
| M F607191 | 2780 | Huang | 4 | 41 | 0.098 | 3 | 0 | 0.750 | 0.000 |
|  |  | Chen | 1 |  | 0.024 | 0 | 0 | 0.000 | 0.000 |
| F15823 | 5190 | Huang | 38 | 2479 | 0.015 | 3 | 11 | 0.079 | 0.289 |
|  |  | Chen | 96 |  | 0.039 | 3 | 23 | 0.031 | 0.240 |
|  |  | Li | 129 |  | 0.052 | 8 | 15 | 0.062 | 0.116 |
| M F15397 | 4980 | Huang | 24 | 1869 | 0.013 | 3 | 21 | 0.125 | 0.875 |
|  |  | Chen | 73 |  | 0.039 | 2 | 21 | 0.027 | 0.288 |
|  |  | Li | 86 |  | 0.046 | 5 | 11 | 0.058 | 0.128 |

The paternal genetic haplotype branch that is specific to the Huang surname in Hubei should exhibit distinct characteristics, with the frequency of Huang surnamed individuals in Hubei being higher within that branch compared to other regions. In each major paternal haplotype branch with at least five Huang samples, and dating back around 4,000 years, we analyzed the proportion of Huang surnamed individuals from Hubei relative to the total number of Huang surnamed individuals nationwide. Branches with a higher proportion of Huang surnamed individuals from Hubei are likely to represent the most Hubei-representative Huang surname branches. These branches are, therefore, more likely to be the ones that originated from the State of Huang and subsequently migrated to Hubei, aligning with the historical records regarding the origins of the Huang surname.

We analyzed the distribution of the Huang surname across various branches downstream of the main paternal haplotype O-F175, which accounts for approximately 75% of the Han Chinese population. These branches include the downstream sub-branches of three major super-grandfather genotypes: M117, F46, and F11 (Table 1 and Supplementary Table S1) (6). Additionally, we analyzed other haplotypes, including those typical of Europeans and Africans (Supplementary Table S1). As a control for the Huang surname, we also analyzed the distribution of the Chen surname, which has a slightly higher population than Huang, particularly in southern China (Table 1).

For the vast majority of haplotype branches, the proportion of the Huang surname within each branch closely aligns with population expectations. The Chen surname also shows similar consistency with expectations. For example, the proportion of Huang surnames within the M117 branch is 0.025, which is close to the expected population proportion of 0.024 for Huang in the dataset.

The Huang surname appears across a wide range of haplotypes common in East Asia, including O, N, Q, C, and D, as well as in some haplotypes common in Europe and Africa, such as R and E. However, it is not found in the majority of non-East Asian haplotypes, including A, B, G, H, I, J, L, M, S, and T (Supplementary Table S1). This broad distribution of the Huang surname across various haplotypes is a common characteristic of large surnames, indicating that most individuals with the Huang surname do not share a closely related common ancestor, or a founder from around 4000 years ago when surnames began to take form.

This observation aligns with the historical records in *Shiji*, which suggest that the citizens of a kingdom naturally include individuals from diverse genetic backgrounds. If these individuals adopted the kingdom’s name as their surname following the fall of the state, the genetic background or paternal haplotypes of Huang citizens would naturally be very diverse. However, there should be a dominant or more common Huang haplotype representing the surname, and individuals from this larger family would be more likely to be from a multi-generational, hereditary royal family. If the citizens of Huang State, along with the royal family, largely migrated to Hubei after the fall of the state, the most common haplotype associated with the Huang surname in the region should exhibit a distinctive high-frequency characteristic in Hubei.

The results indicate a significantly higher-than-expected frequency of Huang surname in Hubei for certain branches. Specifically, in the A16636 branch, the proportion of Huang surname from Hubei is 0.5, which is much higher than the expected proportion of 0.047 (Table 1). Additionally, upstream haplotypes related to A16636, such as A16635, F2137, F2643, F1754, and ACT7481, also show higher-than-expected proportions of Huang surnames from Hubei, with frequencies around 0.14. As a comparison to the Hubei Huang surname, we also analyzed the distribution of Guangdong Huang surname samples in the major branches downstream of M117, as shown in Table 1, but did not find any branches with a notably high proportion of Guangdong Huang samples within the national Huang surname population.

Furthermore, the MF14296 branch, which is downstream of A16636, shows a high proportion of Huang surnames from Hubei as well (0.5), with the proportion of Huang samples among all surname samples within this branch reaching 0.11, significantly higher than the national population ratio of 0.022 (Table 2). Notably, the proportion of Hubei samples among nationwide samples in the MF14296 branch is 0.2, significantly higher than the census result of 0.043 (Table 2), further emphasizing the distinctiveness of this branch in Hubei. These findings strongly suggest that this branch of the Huang surname, A16636-MF14296, has a high-frequency presence in Hubei, aligning with the historical records linking the Huang surname to the region.

**Table 2.** Haplotype distribution for the surnames Huang, Xu, and Liang. Branches with fewer than 5 samples nationwide are not listed. For listed branches that have fewer than 5 samples nationwide in certain surnames, the data are not shown.

| Haplotype | TMRC A | Nationwide | Hubei Huang | Nationwide | Su- Hu Xu | Nationwide | Hebe i Liang | Proportion nationwide |  |  |  |  |  |
| --- | --- | --- | --- | --- | --- | --- | --- | --- | --- | --- | --- | --- | --- |
|  | year | Huang | ratio | Xu | ratio | Liang | ratio | Huang | Xu | Liang | Hubei | Su Hu | He bei |
| Cens us |  |  | 0.042 |  | 0.13 |  |  | 0.022 | 0.014 | 0.007 | 0.043 | 0.076 | 0.054 |
| O- MF14296 | 3580 | 6 | 0.50 |  |  |  |  | 0.11 |  |  | 0.20 |  |  |
| O- MF15137 | 3420 | 6 | 0.50 |  |  |  |  | 0.13 |  |  | 0.21 |  |  |
| O- A16636 | 4320 | 6 | 0.50 | 9 | 0.11 |  |  | 0.008 | 0.012 |  | 0.04 | 0.11 |  |
| C- MF1017 | 3760 | 19 | 0.53 |  |  |  |  | 0.05 |  |  | 0.1 |  |  |
| C- Z45203 | 2980 | 14 | 0.57 |  |  |  |  | 0.09 |  |  | 0.084 |  |  |
| O- MF38096 | 3370 |  |  | 5 | 1.00 |  |  |  | 0.1 |  |  | 0.25 |  |
| O- SK1726 | 2830 |  |  | 5 | 1.00 |  |  |  | 0.13 |  |  | 0.33 |  |
| O- F1442 | 5200 | 13 | 0.08 | 10 | 0.70 |  |  |  | 0.018 |  |  | 0.13 |  |
| O- B432 | 4090 |  |  | 8 | 1.00 |  |  |  | 0.039 |  |  | 0.30 |  |
| O- MF1198 | 4770 | 9 | 0.00 | 14 | 0.57 |  |  |  | 0.018 |  |  | 0.38 |  |
| O- MF941 | 4860 |  |  | 5 | 0.60 |  |  |  | 0.025 |  |  | 0.15 |  |
| O- MF6935 | 3440 | 5 | 0.20 | 7 | 0.57 |  |  |  | 0.012 |  |  | 0.12 |  |
| O- MF6934 | 5330 | 14 | 0.07 | 20 | 0.60 |  |  |  | 0.021 |  |  | 0.17 |  |
| O- MF1075 | 4450 | 18 | 0.11 | 25 | 0.72 |  |  |  | 0.02 |  |  | 0.38 |  |
| O- MF6080 | 4450 | 26 | 0.00 | 6 | 0.67 |  |  |  | 0.011 |  |  | 0.18 |  |
| O- PH4822 | 3460 |  |  | 8 | 0.50 |  |  |  | 0.036 |  |  | 0.19 |  |
| O- CTS5664 | 4570 |  |  | 5 | 0.80 |  |  |  | 0.017 |  |  | 0.087 |  |
| O- Z17557 | 3500 | 11 | 0.00 | 5 | 0.60 |  |  |  | 0.015 |  |  | 0.34 |  |
| O- CTS679 | 3570 | 12 | 0.00 | 6 | 0.50 |  |  |  | 0.012 |  |  | 0.079 |  |
| O- MF18577 | 3960 | 5 | 0.00 | 19 | 0.58 |  |  |  | 0.032 |  |  | 0.42 |  |
| O- MF1107 | 4130 | 17 | 0.12 | 6 | 0.67 |  |  |  | 0.028 |  |  | 0.34 |  |
| O- MF182502 | 4740 | 5 | 0.00 | 23 | 0.61 |  |  |  | 0.033 |  |  | 0.40 |  |
| O- MF9896 | 5020 |  |  | 6 | 1.00 |  |  |  | 0.057 |  |  | 0.12 |  |
| O- Y89945 | 5190 | 6 | 0.00 | 5 | 0.60 |  |  |  | 0.015 |  |  | 0.43 |  |
| O- MF14135 | 5280 | 9 | 0.00 | 41 | 0.54 |  |  |  | 0.04 |  |  | 0.32 |  |
| N- FT317554 | 4480 |  |  | 5 | 0.60 |  |  |  | 0.028 |  |  | 0.19 |  |
| N- F839 | 5050 | 5 | 0.20 | 8 | 0.63 |  |  |  | 0.023 |  |  | 0.23 |  |
| C- F10085 | 4230 | 6 | 0.00 | 5 | 0.60 |  |  |  | 0.0092 |  |  | 0.047 |  |
| C- SK1038 | 4800 | 37 | 0.38 | 8 | 0.50 |  |  |  | 0.007 |  |  | 0.030 |  |
| C- MF10317 | 5320 |  |  | 8 | 0.75 |  |  |  | 0.04 |  |  | 0.22 |  |
| O- MF15764 | 4990 | 6 | 0.17 | 9 | 0.33 | 10 | 0.4 | 0.009 | 0.014 | 0.015 | 0.027 | 0.09 | 0.074 |
| O- MF14287 | 4090 |  |  |  |  | 5 | 0.80 |  |  | 0.1 |  |  | 0.19 |
| O- MF15398 | 3210 |  |  |  |  | 5 | 0.80 |  |  | 0.11 |  |  | 0.2 |
| O- MF14963 | 1850 |  |  |  |  | 5 | 0.80 |  |  | 0.11 |  |  | 0.20 |
| D- F2625 | 5450 | 5 | 0.20 |  |  | 5 | 0.60 |  |  | 0.0082 |  |  | 0.056 |
| O- CTS723 | 3190 | 5 | 0.20 | 5 | 0.40 | 5 | 0.60 |  |  | 0.0073 |  |  | 0.075 |

As shown in Supplementary Table S1, among the 189 branches tested, the vast majority did not exhibit the high-frequency characteristic of the Huang surname in Hubei. Only the C-MF1017 branch and its downstream C-Z45203 branch are exceptions, with frequencies of 0.53 and 0.57, respectively (Table 2). However, the number of Huang samples in the C-MF1017 branch accounts for a very small proportion of all surnames in the country, at only 0.05, and does not show the characteristic of Huang surname population concentration (Table 2). Furthermore, the fraction of Hubei individuals in this branch among all individuals nationwide is merely 0.1, which did not indicate a significant enrichment specific to Hubei.

For the C-Z45203 branch, the Huang surname proportion is relatively higher across all surnames in the country (0.09) (Table 2), but the fraction of Hubei individuals among all individuals nationwide is not high (0.084). Finally, it is worth noting that most Huang surname samples in these two branches come from Huangshi, Hubei. Among the five cities in Hubei with Huang in their names, Huangshi is geographically farther from the Huang state. Therefore, although the C-MF1017 and downstream Z45203 branches show some degree of regional specificity in Hubei, their Hubei specificity is somewhat lacking when compared to the previously mentioned O-MF14296 branch.

As a comparison, we analyzed the proportion of Huang surname samples from Jiangsu in the nationwide Huang surname population (Supplementary Table S1) but found no branches with a high frequency of Jiangsu Huang surnames (= or > 0.5). Although one branch, O-F4068, has a relatively high frequency (0.46), the proportion of Huang surnames within all surnames in this branch is only 0.026 (Supplementary Table S1), which is similar to the census proportion. This suggests that the presence of a high-frequency Huang surname branch specific to Hubei indicates a certain level of regional specificity.

Before analyzing the 23Mofang database, we had previously conducted high-throughput sequencing on an individual with the Huang surname from Hubei and uploaded the data to the 23Mofang database. The terminal genetic marker for this individual was MF287231, a downstream marker of A16636 (Figure 1). His paternal family resides in Huangjiadun Village, Luohansi, Huangpi District, Wuhan, where Huang is a prominent surname. According to the family genealogy, other closely related Huang surname families in the area include eight villages named after Huang, such as Huangbafwan, Huangjiadawan, Huangjiagang, Huangjiadang, Huangjiayuanzi, Huangjiadaduwu, Huangjiaxiaoduwu, and Huanghuxiwan (7).

**Figure 1:**
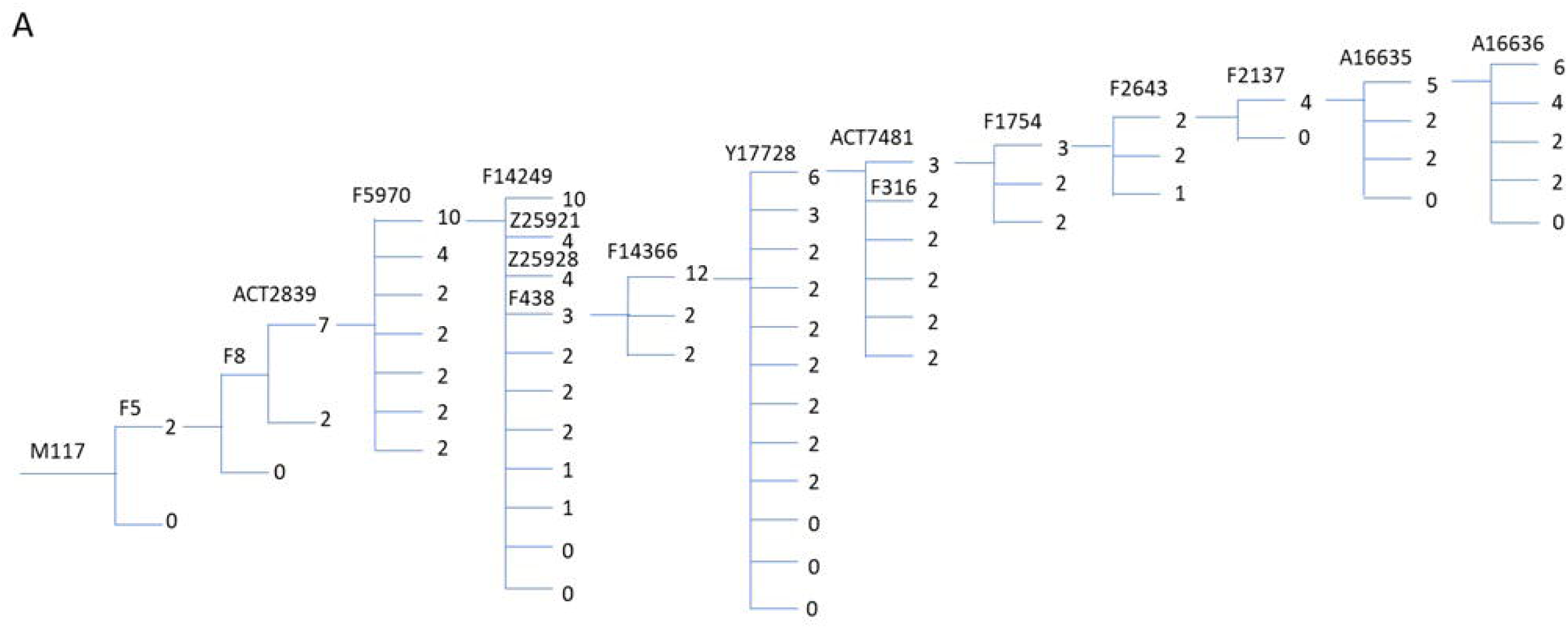
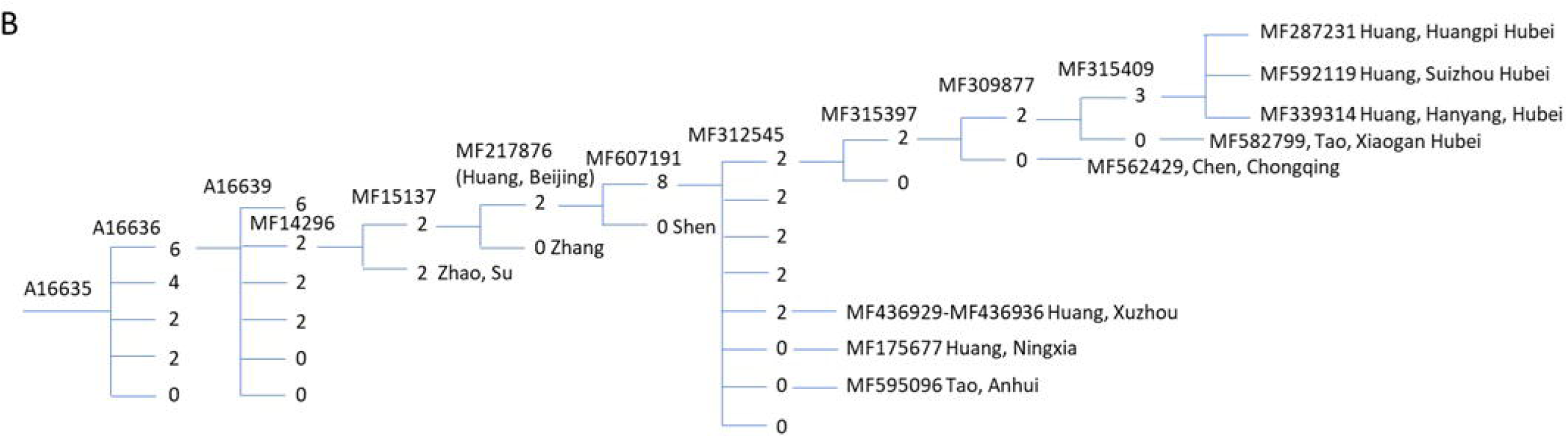
Paternal lineage phylogenetic tree of a Huang surname Individual (MF287231) from the Huangpi Luohansi Huang Family in Hubei. The tree is from 23Mofang. The genetic markers of the selected branches are shown above the branch lines, with the number of descendant branches for each lineage also indicated in the figure. Sister lineages are arranged from top to bottom according to the number of descendant branches, from most to least. A. Phylogenetic tree from M117 to A16636. B. Phylogenetic tree from A16635 to MF287231.

To verify if the mainstream genetic marker of these villagers is A16636, we conducted sampling and sequencing experiments. We randomly sampled seven Huang surname individuals, including three from Huangjiadun Village, two from Huangjiadawan Village, and two from a neighboring village, Lianglukou, which does not have "Huang" in its name and is not related to the family tree. The results showed that, out of the three individuals from Huangjiadun Village, two carried the A16636 marker, while one did not; further investigation revealed that the third individual was adopted. Both individuals from Huangjiadawan Village carried the A16636 marker, while the two individuals from Lianglukou Village did not. These findings suggest that the Huang surname families from the nine villages represented by Huangjiadun Village should all share the A16636 genotype.

In the 23Mofang database, there are six Huang surname samples nationwide with the A16636 marker (seven samples were displayed, but two from Ningxia are father and son, so only one was counted). Three of them did not have downstream branch sequencing information, which we sponsored for deep sequencing (only the father was sequenced in the Ningxia father-son pair). The results showed that all six individuals belong to the MF14296 branch.

The two branches closest to the MF287231 individual are a Huang surname individual from Han Yang District, Wuhan (MF339314), and a Huang surname individual from Suizhou, Hubei (MF592119). Han Yang is located 40 kilometers south of Huangpi, and Suizhou is located 80 kilometers northwest of Huangpi. The most recent common ancestor haplotype for these three individuals is MF315409, with a common ancestor time estimate ranging from 510 to 1210 years ago as calculated by 23Mofang (Figure 1). These two family branches do not belong to known relatives from the Huangjiadun Village. This result suggests that there might be a Huang surnamed ancestor individual living in the northeastern part of Hubei during the Ming, Yuan, or Song dynasties (510– 1210 years ago).

### The estimated origin date for the Hubei Huang surname

The naming ancestor of the Huang surname branches downstream of A16636 in Hubei should likely be traced to a specific period or branch. Given that three Hubei Huang surname individuals carry the MF315409 marker, with a common ancestor time estimated between 510-1260 years ago, it can be inferred that they likely had a Huang surname ancestor living in Hubei at that time. The upstream branch, MF309877, has a common ancestor time between 1260-2080 years ago, and a Hubei Tao surname individual (MF582799) also carries this marker (Figure 1). This suggests that there was an individual in Hubei, likely carrying the MF309877 marker, whose descendants remained in Hubei. Most of them were Huang surname descendants, but a small portion of them later adopted the Tao surname. Consistent with this inference, A16636 Hubei Tao surname individuals account for only 0.25 of the total Tao surname population in the same branch, without showing a high concentration in Hubei (whereas A16636 Hubei Huang surname individuals account for 50% of the total Huang surname population in the same branch).

The Huang surname branch closest to the MF309877 branch is a Huang surname branch from Xuzhou, which belongs to the MF436936 branch. The common ancestral marker between these two different Huang surname branches is MF607191, with a common ancestor time estimated between 2950-3310 years ago. This branch has 41 individuals, with 11 (0.268) from Hubei, and the highest number from other provinces is only 4 individuals. This suggests that the population of this branch is more concentrated in Hubei, indicating that the ancestors likely lived in Hubei. Furthermore, because the proportion of Huang surname individuals among the Hubei descendants of this branch is relatively high (0.27), it can be inferred that the ancestor of this branch likely carried the Huang surname. The other surnames among the Hubei descendants may have resulted from surname changes or other factors.

MF217876 is the closest upstream branch to MF607191, with a common ancestor time between 3310-3420 years ago. A Huang surname individual from Beijing carries the MF217876 marker but does not carry MF607191. This branch only has 3 more individuals than the MF607191 branch, but it includes 1 Huang surname individual. This suggests that the ancestor of the individuals carrying the MF217876 marker likely had the Huang surname, and some of the Huang surname descendants migrated to other regions.

MF15137 is the closest upstream branch to MF217876, with a common ancestor time between 3420-3610 years ago. It only has 2 more individuals than MF217876. Therefore, it is highly likely that the Hubei individuals carrying MF15137 have ancestors from Hubei, and their ancestor most likely carried the Huang surname.

MF14296 is the closest upstream branch to MF15137 and the closest downstream branch to A16636, with 10 more individuals than MF15137 (total 56 individuals), and 11 of them are from Hubei (0.20), the highest proportion in the country. Other provinces with relatively high representation include Jiangsu (5 individuals), Shandong (5 individuals), Hebei (5 individuals), Zhejiang (4 individuals), Anhui (3 individuals), and Henan (3 individuals). The proportion of Huang surnames (6 individuals) in this branch compared to all surnames in the same branch is 0.11, which further suggests that the Hubei ancestors of the individuals carrying MF14296 likely carried the Huang surname. Other prominent surnames in the MF14296 branch with a significant number of individuals include Zhang (5 individuals), Ren (5 individuals), and Li (4 individuals). In contrast, A16639, a sister branch of MF14296, does not exhibit the same high proportion from Hubei (0.018), further supporting the Hubei specificity of the MF14296 branch.

The proportion of individuals from Hubei carrying A16636 is 0.042, and the Huang surname in this branch accounts for only 0.007 of all surnames in the same branch, which is three times smaller than the expected population proportion. This indicates that the majority of descendants carrying A16636 do not exhibit a concentration in Hubei, nor is there a concentration of the Huang surname. Therefore, it is highly likely that the ancestor carrying A16636, from which the Hubei Huang surname branch of MF14296 originates, was not originally a Huang surname and did not live in Hubei.

Based on the above analysis, the earliest ancestral marker for the Huang surname in Hubei, located downstream of A16636, is likely MF14296, but MF15137 is also possible. Based on these two markers, a common ancestor can be estimated to have lived between 3420 and 4320 years ago, corresponding to the early or middle period of the Huang state’s history. This suggests that the Huang surname likely originated before the fall of the Huang state and could only have been a surname belonging to the royal family of the Huang state. Therefore, it is possible that the royal lineage of the Huang state is associated with the MF14296 or MF15137 branch.

### Analysis of the Xu surname

If the historical genetic lineage of the Huang surname aligns with the historical records such as *Shiji*, which state that it originates from the eldest son of Bo Yi, named Da Lian, then the Xu surname, being descended from the second son of Bo Yi, named Ruo Mu, is the most likely to possess a similar genetic history that aligns with the historical records. The Xu surname, originating from the people of the Xu State, adopted the name of their state and became concentrated in Jiangsu. Therefore, the founding genetic branch of the Xu surname should exhibit a high-frequency, region-specific feature in Jiangsu.

Similar to the study conducted on the Huang surname, we analyzed the 189 haplotype branches and calculated the proportion of the Xu surname in Jiangsu and Shanghai (Su-Hu) among the same surname population nationwide within the same branch (Table 2 and Supplementary Table S1). We found that the Xu surname in Su-Hu is concentrated in the branch A16635-F15823-F1442-MF38096 (with a common ancestor from 3370-5200 years ago). This branch has 100% of the Xu surname population in Su-Hu. It is also enriched with the Xu surname among all surnames nationwide with a significant proportion of Xu (0.104), much higher than the national population ratio of Xu (0.016). Furthermore, the proportion of Su-Hu individuals among all individuals nationwide in this branch is also relatively high (0.25).

We also identified 23 other branches, where the proportion of Xu surname from Su-Hu within the national Xu surname population is relatively high (greater than 0.5, Table 2 and Supplementary Table S1). However, when considering the proportion of the Xu surname in the total population of all surnames, these branches are at relatively low levels (0.008-0.057), and therefore do not belong to branches where the Xu surname is notably concentrated. Although some of these branches have characteristics of high frequency of Su-Hu population, they do not show a concentration of Xu surname population. For example, in branch O-Y89945, the Su-Hu population accounts for as much as 0.43, but the proportion of Xu surname in this branch is only 0.015. Therefore, compared to branch O-MF38096, these branches do not represent the most Su-Hu specific Xu branch.

Although the upstream branch F1442 of MF38096 shows a high proportion of the Xu surname in the populations of Jiangsu and Shanghai (0.7), it does not exhibit the characteristic of a high concentration of Xu surname within all surnames (only 0.018) (Table 2). The Xu surname samples from MF38096 are concentrated in its downstream branch SK1726 (common ancestor time 2830-3480 years ago) (Table 2), but these Xu surname samples lack information from downstream branches of SK1726. The sample size of SK1726 is 40 people, accounting for 73% of the total samples from MF38096. Both of these branches generally meet the standards for the ancestral branch of the Xu surname, but the upstream F1442 branch of MF38096 clearly does not. Therefore, the ancestral branch of the Xu surname is likely MF38096, but it cannot be ruled out that it could be SK1726. This is generally consistent with historical records of the State of Xu and the origin of the Xu surname. This Xu surname branch shares a recent common ancestor with the Huang surname branch identified earlier, which is A16635, further confirming that the Huang surname originates from the Ying surname as recorded in *Shiji*.

### Analysis of the Liang surname

The Liang surname belongs to the descendants of Fei Zi, a distant son of the Bo Yi family. The state of Liang was established around 800 BCE, and after its fall, the population adopted the surname Liang and migrated to the state of Jin, which is now in the Shanxi and Hebei regions. Similar to the analysis conducted above, we studied a total of 189 branches and analyzed the proportion of Liang surnames from Shanxi or Hebei within the national Liang surname population.

The results revealed that five branches showed a high proportion of Liang surname in Hebei (greater than 0.5) among the national Liang surname population (Table 2). Among them, only the MF14287 branch (4090-4900 years ago) and its two downstream branches, MF15398 and MF14963, had a significantly higher proportion of the Liang surname within all surnames (0.1), much higher than the general Liang surname proportion of 0.007. However, MF15764, which is the upstream branch of MF14287, showed a Liang surname proportion of only 0.015 in all surnames (Table 2). Further analysis revealed that the Liang surname samples are all within the MF14963 branch (common ancestor time 1850-3210 years ago), with a proportion of 0.1 within all surnames in this branch. Additionally, this branch has the highest proportion of Liang surname population in Hebei (0.21), and at least two different downstream branches of this branch contain Liang surname samples. These results suggest that the ancestral branch of the Liang surname from the state of Liang is most likely MF14963, but it cannot be ruled out that it could also be MF14287 or MF15398.

We did not find any branch with a high proportion of Shanxi Liang surnames within the national Liang surname population (Supplementary Table S1), suggesting that the Liang surname group from the state of Liang, which belonged to the MF14287/MF15398/MF14963 branch, likely passed through Shanxi and migrated to Hebei. This also implies that there is no branch in Shanxi where the Liang surname is significantly more concentrated compared to the national population. Overall, the finding of the Liang surname aligning with the historical records further supports the genetic conclusions drawn from the Huang and Xu surnames. The consistency observed in the study of different surnames further strengthens the argument that genetic data can provide valuable insights into the historical origins of these families.

The present-day individuals sharing the most recent ancestor branch with the Liang surname, MF14287, include the Zuo family from Xiangyin, Hunan, whose genetic marker is a more downstream branch, MF59804 (Figure 2). Among the 23Mofang users in this branch, there are 5 individuals, excluding 1 close relative, leaving 4 effective individuals, 75% of whom are from the Zuo surname in Hunan. As one of the prominent officials of the late Qing dynasty, the national hero Zuo Zongtang’s ancestral home is in Xiangyin, Hunan, so he is very likely to belong to this branch.

**Figure 2:**
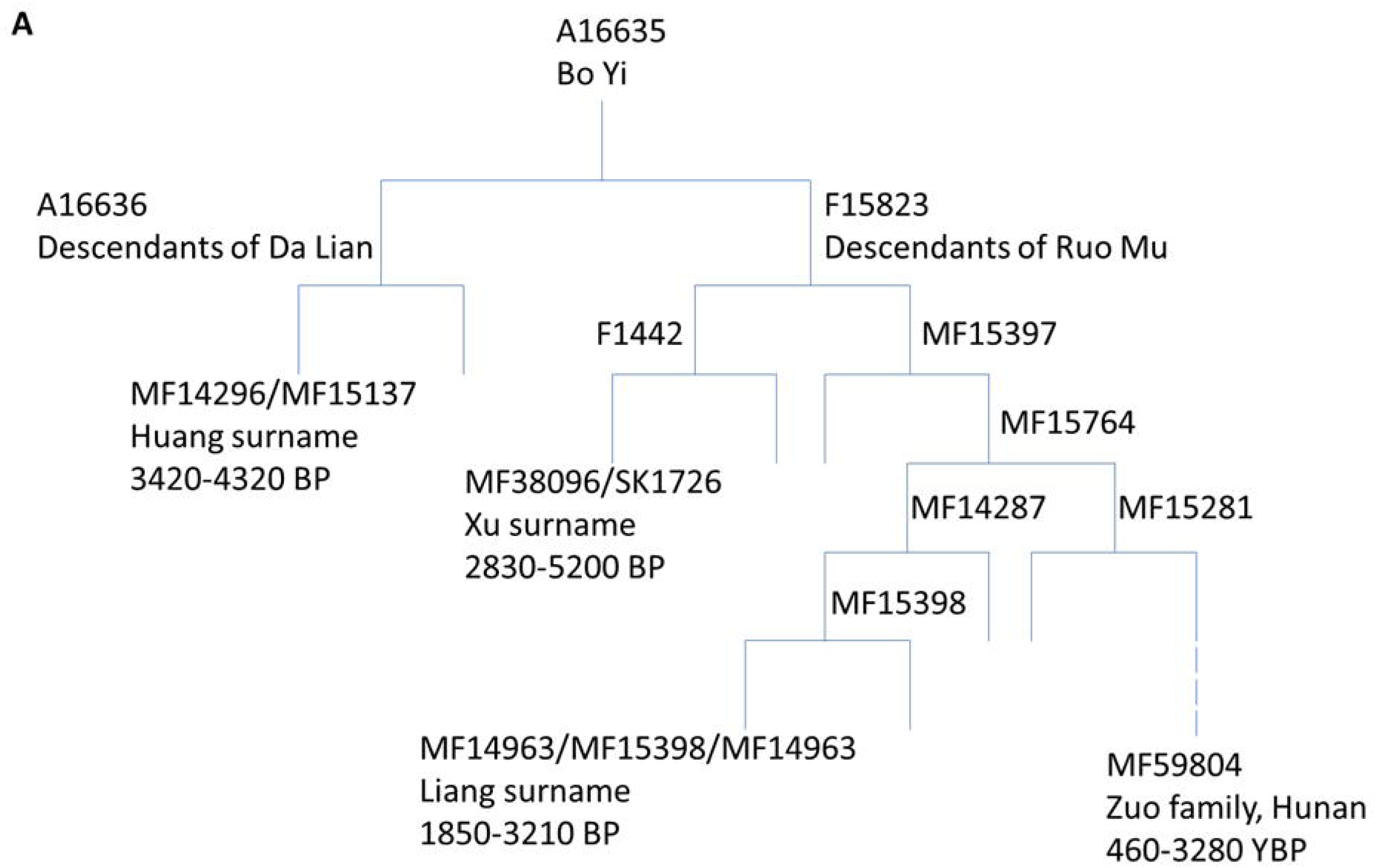
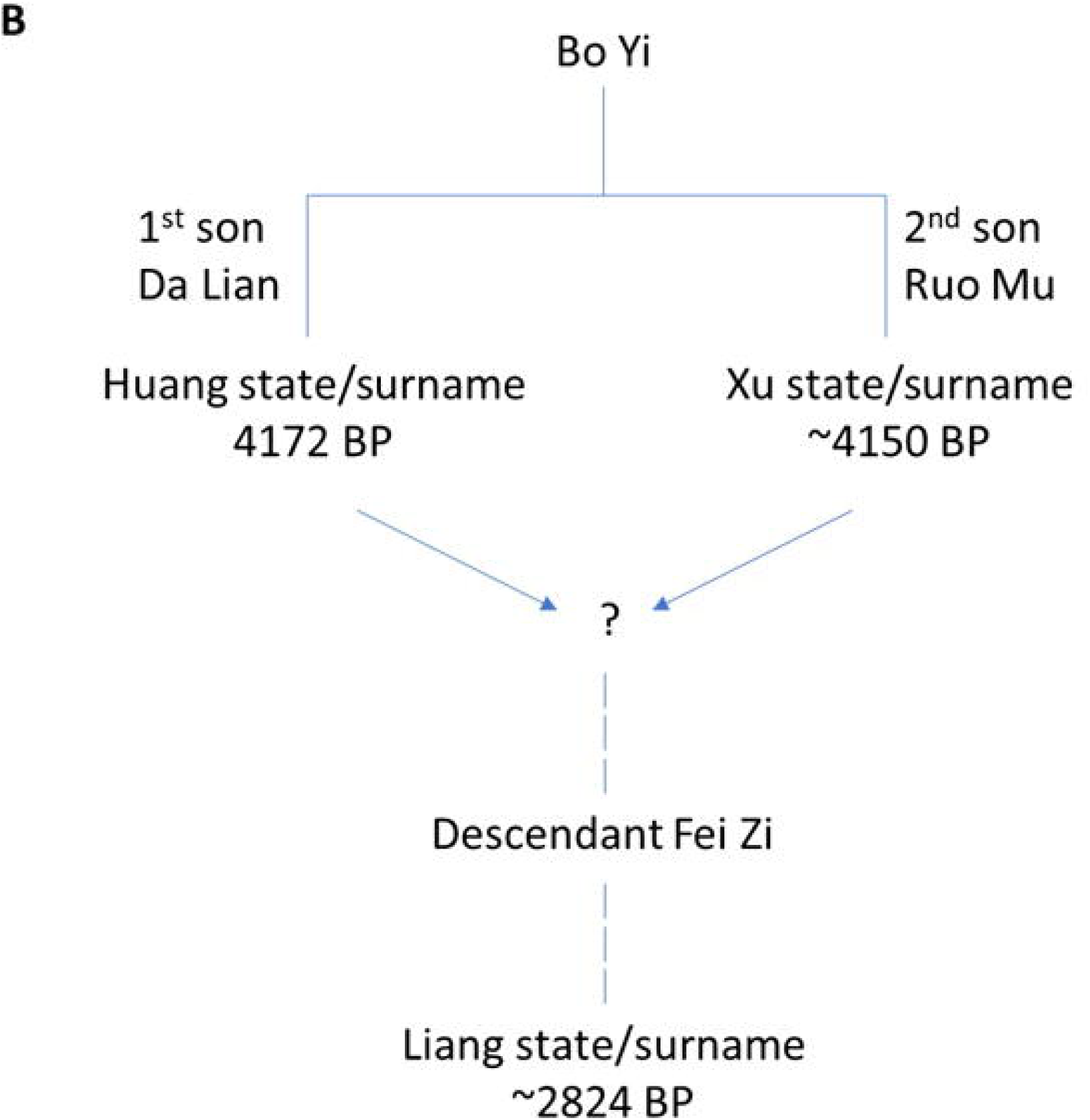
The family tree of surnames related to the Ying surname. A. Phylogenetic tree of Bo Yi and his male descendants. BP: before present. B. The genealogy of the Bo Yi clan, as recorded in historical texts such as *Shiji*.

### Other surnames under Ying

We also analyzed the genetic distribution of two other more populous Ying surnames, Zhao and Ma, but did not find any high-frequency genetic branches that align with the historical records of their settled regions. This may be due to these surnames being more distantly related to Bo Yi. The ancestors of the Zhao and Ma surnames are both attributed to Zao Fu, who is recorded as the 14th descendant of Bo Yi. Additionally, the more recent Zhao emperor, Zhao Kuangyin from the Song Dynasty, likely had a large number of descendants, which may have caused changes or confusion in the Zhao family tree. Other less populous Ying surnames cannot be meaningfully analyzed at this time and will require more data from future 23Mofang customers for further study. Furthermore, as these surnames are more distantly related to Bo Yi or have less certain historical connections, they are of limited use in verifying the historical records.

### Other Region-Specific Huang Surname Lineages

To gain a comprehensive understanding of the regional specificity of the paternal genetic lineages associated with the Huang surname, we further analyzed the proportion of the Huang surname within each of the 189 lineages mentioned above (Supplementary Table S1). Most lineages showed a proportion consistent with census data, which is 0.022. Apart from the aforementioned Hubei-specific lineages MF14296/MF15137 (with proportions of 0.11 and 0.13, as shown in Table 2), only three other lineages showed a relatively high proportion (ranging from 0.09 to 0.28), with clear regional characteristics (the proportion of the Huang surname samples in these regions was higher than 0.5 among all Huang surname samples), including O-MF35829, O-Z25482, and O-F1094 (Table 3). Among these three lineages, the first two are Guangdong specific and the third one is Zhejiang specific. However, these lineages did not show close relationships with the Xu surname lineages of the Su-Hu regions as documented in the literature. The Guangdong Huang surname lineage of MF35289 was concentrated in its downstream branch Y28920 (formation time 2270 years ago), and this Huang surname lineage has a sister branch, MF9896 (formation time 5020 years ago). The two sister branches share an upstream branch, MF9858 (formation time 9460 years ago). Although the MF9896 lineage is enriched with the Xu surname samples in the Su-Hu region, the proportion of samples from the Su-Hu region among nationwide samples is not high (0.12). Also, the shared ancestor time (9460 years ago) is significantly earlier than the era of the Bo Yi. Additionally, Guangdong is geographically distant from the state of Huang. Therefore, this Guangdong Huang surname lineage MF35289 does not align with the narrative in historical records regarding Da Lian and Ruo Mu.

**Table 3.**
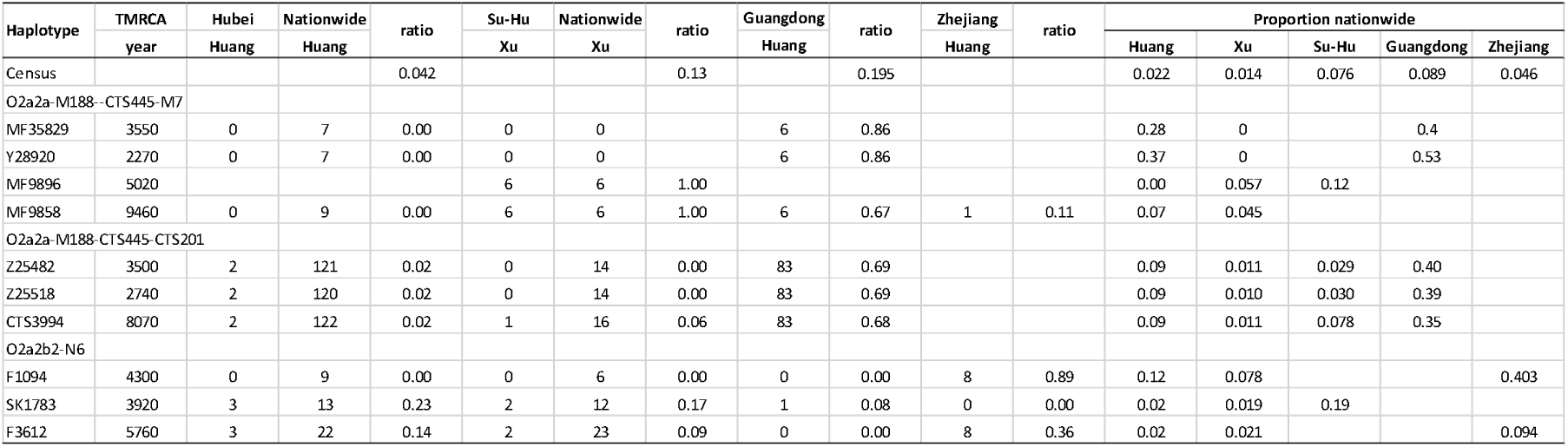
The proportion of Huang and Xu surnames in different haplotypes and regions. For listed branches that have fewer than 5 samples nationwide in certain surnames, the data are not shown.

The Guangdong-specific Huang surname lineage also includes O-Z25482, but this lineage does not have a sister branch concentrated with the Xu surname in the Su-Hu region, thus it does not align with the historical records.

The Zhejiang-specific Huang surname lineage F1094 has a sister branch Y170907-SK1783, and they share the ancestral marker F3612. However, the SK1783 branch is not enriched with Xu surname samples, and the proportion of Xu surname nationwide is also not high (0.019). Therefore, the Zhejiang Huang surname lineage F1094 does not align with the historical records. Overall, while there are several region-specific Huang surname lineages outside Hubei, these lineages do not align with the historical narratives regarding the states of Huang and Xu. This also indicates that the Hubei-specific Huang surname lineage MF14296/MF15137 is the only one among all tested Huang surname lineages that aligns with the historical records.

## Discussion

This study utilized the 23Mofang genetic database in combination with independent and sponsored sequencing studies to systematically analyze the distribution of the Huang surname and other Ying-related surnames across different paternal haplogroup branches. The study was inspired by the *Shiji* and other historical literature and arrived at genetic conclusions that align closely with the historical records in *Shiji*.

Based on literature and the current population distribution of the Huang surname, the Huang populations most closely associated with the origin of the surname should primarily be located in Hubei. The analysis in this study suggests that these populations may trace their ancestry to a Huang family of the Huang Kingdom, with a genetic marker MF14296/MF15137, dating back approximately 3,420-4320 years.

If the historical records such as *Shiji* regarding the origin of the Huang surname are accurate, the following conclusions should hold:

1. The Huang Kingdom should have existed, and the Hubei Huang surname group, originating from the Huang Kingdom, is most likely the group that best matches the recorded Huang surname population. The formation of the Huang surname should not date later than the fall of the Huang Kingdom, which occurred 2,672 years ago.
2. Figures from the Eastern Yi families, such as Bo Yi, and Da Lian, should be historically accurate. These ancient Huang Yi tribal figures, along with their ancestors like Huang Di, should exhibit traits of super-grandfathers, meaning their paternal lineage should have many descendant branches.
3. The Huang ancestor family should belong to the Xia Dynasty, and since the Huang state was also founded during the Xia Dynasty, the Xia Dynasty should have existed.
4. Other clearly documented surnames belonging to the Ying family, such as the Xu and Liang surnames, should also be valid and genetically related to the Huang surname lineage.

The existence of the Huang state is supported by archaeological evidence, located in Huangchuan, Henan. However, the exact starting time of the kingdom’s existence in relation to the Xia Dynasty has not yet been confirmed through archaeological investigation (8). This study finds that the formation time of the initial Huang surname branches dates back to 3,420–4,320 years ago, corresponding to the early or middle period of the Huang state. The discovery of a Huang surname branch, MF14296/MF15137, in Hubei, which aligns with historical records, is likely not a coincidence, but rather reflects a certain degree of objectivity in the historical texts. As a comparison, we were unable to make similar discoveries in an unrelated province such as Jiangsu.

Although our research discovered a C-MF1017 branch and its downstream branch C-Z45203 within the Huang surname population in Hubei, which also show a relatively high proportion of Huang surnames in the national Huang population, these branches are not as consistent with the characteristics that should be associated with the ancestral origin of the Huang surname, as seen in the O-MF14296 branch. This includes a lower proportion of the Huang surname in the total population of surnames nationwide, the fact that most Huang samples come from areas farther from the ancient Huang state, and a relatively low proportion of samples from Hubei in these branches compared to the national total.

Due to the significant impact of surname changes or ancestral recognitions (such as granted surnames), the haplotype of an ancient surname cannot be definitively determined by its national proportion within the corresponding haplotype. Surnames and their associated haplotypes typically exhibit regional specificity. The connection between Hubei Huang surnames and the MF14296/MF15137 haplotype is indicative of the regional characteristics of surname haplotypes. This adds complexity to the study of surname gene branches. It cannot be simply inferred from national proportions alone. Furthermore, it highlights the importance of conducting regional-specific research on the attributes of a surname gene branch, particularly for ancient surnames that have been in use for thousands of years. If a surname and its associated haplotype originate from a specific region or province, that region’s population should have the highest proportion of that surname and haplotype compared to the rest of the country.

The Huang Clan Kinship Network (http://www.ihuang.org/a4.htm) lists many haplotypes associated with specific Huang surname families. Our study similarly found that the Huang surname is also concentrated in many other haplotypes. However, these haplotypes do not align with the historical records of the Huang surname’s origin. From the perspectives of time and location, only the Hubei Huang haplogroup MF14296/MF15137 most closely matches the historical and archaeological records of the Huang State and the Huang surname. Therefore, the different genetic branches of Huang surname populations likely do not stem from a recent common ancestor with the surname Huang, but rather, these individuals may have independently adopted the surname Huang.

Various complex factors, such as surname changes, granted surnames, adoptions, and the recognition of ancestral roots, can obscure the traces of the surname’s origin. While the Huang surname is present in nearly all major branches, it is clear that only one of these branches represents the Huang surname of ruling family of the Huang State. Combining research on surname and genetic regional specificity with the verification of historical documents could be a more effective new approach in surname genetics.

This study independently supports the four conclusions derived from historical literature from a molecular genetics perspective. The ancestral genotype of the Hubei Huang surname branch MF14296/MF15137 consists of F2137, F8, and M117, which belong to one of the three major super-grandfather genotypes (Figure 1A) (6). This genotype is also the one most closely aligned with the historical records of the Huang Yi tribe and Huang Di (5).

The different ancestral branches of MF14296/MF15137 exhibit characteristics of super-grandfathers. For example, the most recent upstream ancestor, A16636, has six downstream branches, more than any of the other four sister branches at the same level (Figure 1B). Similarly, A16636’s upstream ancestor, A16635, has five downstream branches, again surpassing other sister branches at the same level.

In total, among the 13 ancestral branches from MF14296 to M117, only one branch, F438, does not have the super-grandfather characteristics compared to its sister branches. F438 contains three downstream branches, fewer than F14249 (with ten downstream branches), Z25921 (with four downstream branches), and Z25928 (with four downstream branches). However, F438 has the largest number of descendants, 2.91 times more than F14249, 2.8 times more than Z25921, and 4.2 times more than Z25928. This suggests that the ancestors of the MF14296 Huang surname branch at different stages of time predominantly exhibit characteristics of emperors or noble families, which aligns well with the historical records that describe the Huang surname’s ancestor as the prominent minister Bo Yi of the Xia Dynasty, and traces Bo Yi’s lineage back to Huang Di and the Huang Yi tribe.

At the same time, this study further supports the possibility that the F8/M117 haplotype may belong to Huang Di, as the ancestors of the Huang surname are recorded in historical texts as originating from the ancient Huang Yi tribe, whose leader was titled Huang Di.

A16635 is the primary branch of the descendants of F2137, with 73.2% of the current F2137 descendant population belonging to A16635. Ancient DNA studies show that F2137 was a common haplotype in the Central Plains and surrounding regions around 4,000 years ago. These sites include Mogou in Gansu (4,000 years ago) (9), Shimao in Shaanxi (4,000 years ago) (10), Mianchi Yangshao Village in Henan (4,000 years ago) (11), and Dashanqian in Chifeng of Inner Mongolia (3,500 years ago) (9). This further supports the hypothesis that F2137 is a super-grandfather and aligns with the historical records that describe the Huang and Ying surname ancestors as noble figures.

This study shows that other descendants of Bo Yi, such as the Xu and Liang surnames, which have more clearly documented historical records, also exhibit regional haplotype enrichment similar to that of the Huang surname in Hubei. The founding haplotype of the Xu surname from the state of Xu is likely MF38096/SK1726, which is more frequently found in Jiangsu and Shanghai, while the founding haplotype of the Liang surname from the state of Liang is likely MF14963/MF15398/MF14963, which is more common in Hebei. Their genetic types and the time of appearance of their common ancestors align with historical records such as *Shiji*. The genetic evolutionary relationships between these surnames and their connection to the Huang surname in Hubei also align with historical documentation (Figure 2). Additionally, both the Xu and Liang surnames, like the Huang surname, appear in multiple other haplotypes with significant populations, but these branches do not represent the most ancestral characteristics of the Xu or Liang surnames. Therefore, the three major surnames under the Ying clan, including the two with the clearest historical records and closest ties to Bo Yi—the Huang and Xu surnames—both support the narrative of *Shiji* at the genetic and regional distribution levels.

The genetic characteristics of other Ying surname variants still require further in-depth research to determine. The Zhao surname and Ma surname are recorded as having a common ancestor, Zao Fu, who is a descendant of Bo Yi but further removed. This study was unable to identify the ancestral genotypes of these two surnames. This may be due to their more distant or uncertain connection to Bo Yi, and it could also be related to the influence of the Zhao surname during more recent times, such as Emperor Zhao Kuangyin of the Song Dynasty. The imperial influence, along with practices like surname changes or bestowed surnames, could dilute the ancient lineage characteristics of these surnames. In feudal society, name avoidance was a common practice. People would change their surnames or names to avoid having the same or similar names as the emperor or elders. Additionally, after the fall of the Song Dynasty, some people with the Zhao surname likely changed their surname to avoid persecution under the Yuan Dynasty.

The other surnames of the 14 branches of the Ying clan mentioned in *Shiji* have not been included in this study due to their small population. Moreover, their connection to Bo Yi is less clear in ancient texts, and their correspondence with modern surnames is also less well-defined. Therefore, the study of these surnames has limited significance for the independent verification of the historical records.

If the Huang surname originally came from the citizens of the State of Huang, who took the state’s name as their surname, and the ruling family of Huang also carried a specific haplotype X, we would expect that the majority of the Huang citizens or people with the Huang surname would not carry haplotype X. This is because, even though the descendants of the ruling family would likely outnumber the ordinary families, the population of Huang state had already been substantial when the state was established, and the paternal lineage distribution in the population should reflect the diversity of paternal genetic markers present in the central plains at the time. Additionally, it is possible that a large number of people from other regions joined the State of Huang, which would also increase the genetic diversity of its population. The tradition of adopting the state name as a surname would result in a large number of people, with different genetic backgrounds, sharing the same surname. This would mean that, despite the presence of the ruling family’s haplotype X, the broader population would exhibit a greater diversity in their paternal lineage, and not all Huang surname bearers would share the haplotype X associated with the ruling family.

The founding lineages of the Huang and Xu surnames belong to two distinct branches, A16636 and F15823, both of which are the most recent downstream branches of A16635. A simple and intuitive logic might place Bo Yi under A16635, with his two sons belonging to A16636 (Da Lian) and F15823 (Ruo Mu) respectively. However, while this relationship is not impossible, there could be other explanations. The mutation rate of the Y-chromosome is approximately once every 84.5 years (according to the standard from 23Mofang), so Bo Yi and his sons may have identical Y-chromosomes, but it is also possible that they could differ. If they were the same, it could mean that all three (Bo Yi, Da Lian, and Ruo Mu) were part of the A16635 lineage, with mutations like A16636 potentially occurring in one of Da Lian’s descendants, and F15823 occurring in one of Ruo Mu’s descendants.

The common ancestor time of A16635 (5340-5360 years) significantly differs from that of its downstream branches A16636 and F15823 (4330-5340 years and 5230-5340 years, respectively). This suggests that there might be a nearly 1000-year gap between A16635 and A16636, which indicates that the relationship between A16635 and the Da Lian lineage may not be a direct father-son relationship. This supports the idea that the mutation of A16636 likely did not occur in Bo Yi’s son Dalian, but rather in a more distant descendant of Dalian. However, it is important to note that this 1000-year difference may not be entirely accurate, and the actual difference could be much smaller.

Recent ancient DNA studies and full Y-chromosome sequencing have established a new idea and phenomenon: many mutations previously thought to be ancestral mutations are actually independent convergent mutations that occurred in different downstream branches (12, 13). Therefore, the older an ancestral lineage is, the more it will contain convergent mutations from later generations. These mutations naturally increase the apparent time difference between an ancestor and its closest downstream branches. Given that a mutation occurs on average every 84.5 years, the time difference between an ancestral lineage and its closest downstream lineage should typically be within a few hundred years, not a full 1000 years.

This discrepancy in estimated time suggests that the method of calculating common ancestor time based solely on mutation counts has its limitations and does not account for repeated mutations. This could explain why two branches at the same level, such as A16636 with multiple mutations (9 mutations based on ISOGG or International Society of Genetic Genealogy data website) or a greater temporal gap from A16635, and F15823 with fewer mutations (1 mutation) or closer to A16635, exhibit significant variation in their mutation profiles. This further supports the recent discovery that repeated or convergent mutations are very common in Y-chromosome.

Therefore, the current estimate of the common ancestor time for A16635 by 23Mofang only serves as an approximation. The actual time could be smaller by hundreds of years, or even up to a thousand years. Therefore, the estimated common ancestor times of major lineages like A16635 are not in significant conflict with the historical figures mentioned in *Shiji*, which date to around 4000-5000 years ago. Likewise, the time estimate of the appearance time for F8 (7290-12910 years ago) as calculated by 23Mofang is also merely an approximation with large error margins. And it significantly differs from the time estimated by other researchers (around 5,400 years) (6).

The *Shiji* mentions: "Da Lian’s great-grandson was named Meng Xi and Zhong Yan. He had the body of a bird but spoke human language. The Emperor Tai Wu heard of this and divined it. The result was auspicious, so he appointed Meng Xi to serve as a charioteer and married his daughter to him. From Tai Wu onward, many of Zhong Yan’s descendants had assisted the Shang dynasty with achievements. So, individuals with the Ying surname were commonly prominent (Ying Xing Duo Xian), and became lords." This suggests that Da Lian’s descendants held a high status and influence during the Shang dynasty, and it also supports the idea that Da Lian indeed had male descendants, aligning with the historical records of the Huang State’s royal family, where the lineage passed down for multiple generations over a thousand years.

The phrase "individuals with the Ying surname were commonly prominent (Ying Xing Duo Xian)" being noted by historians and written into historical texts likely reflects its unusual significance. This study suggests that this prominence may be related to genetic factors. Not only was this true during the Shang Dynasty, as the historical records describe, but it might have been the case in ancient times as well. The upstream genetic branches of the Ying surname generally exhibit characteristics of a super-grandfather, such as the most likely branch associated with Huang Di, M117-F8-F2137. The central figure in the unification of China, Emperor Qin Shi Huang, was named Ying Zheng. He is possibly a descendant of Bo Yi, but the specific line of descent is not clearly documented in historical records and requires further in-depth research to determine.

Additionally, in more recent times, descendants of the Ying surname may have also produced notable figures. By identifying the genetic signature of the Ying surname’s ancestors, it becomes possible to link it to famous historical figures. The Zuo family from Xiangyin, Hunan, for example, carries this genetic signature (Figure 2). The late national hero Zuo Zongtang is likely to have originated from this family, although further research is needed to confirm this.

The M117-F8 lineage, over approximately a few thousand years, has grown to now represent about 17% of the Han male population, ranking it first in terms of the number of descendants among the super-grandfathers of the Neolithic age (6). This phenomenon supports the notion of a fitness advantage of this lineage. However, whether there are specific genes on the Y chromosome directly linked to fitness remains to be determined through future, more in-depth research. Existing genetic studies on complex traits such as the GWAS (genome wide association study) type of research have typically omitted genetic variants from the Y chromosome.

The genetic data from this study suggest that Da Lian himself, as well as his immediate descendants, who may have carried the A16635 or A16636 variant, may not have formally adopted the Huang surname. Instead, the formal use of the Huang surname may have originated from a descendant of Da Lian who carried the MF14296/MF15137 marker (Figure 2). This change likely occurred several centuries after Dalian established the Huang state. The *Shiji* notes that the fifth-generation descendants of Dalian had names, Meng Xi and Zhong Yan, that has no apparent link with the Huang surname. Similarly, the formal adoption of the Xu surname did not directly come from Ruo Mu himself or his immediate descendants, but rather from a descendant of Ruo Mu who carried the MF38096 or SK1726 genetic marker (Figure 2). This change likely occurred several centuries after Ruo Mu founded the Xu state.

Previously, it was suggested that the formal and widespread use of surnames began during the Zhou Dynasty (1046–256 BCE), when the feudal system played a key role in the spread of surnames. Although the early forms of surnames existed prior to the Zhou Dynasty, their usage may not have been widespread (8). The findings of this study do not significantly conflict with this view. In fact, they suggest that the formal adoption of the Huang and Xu surnames may have occurred slightly earlier than the Zhou Dynasty, but still within the larger context of the gradual development of the patriarchal system, which influenced social structures. This discovery further enriches our understanding of the origins and evolution of surnames.

Regarding the origin of the Huang surname, there are different historical accounts. One prominent view holds that Lu Zhong, a descendant of Zhuan Xu (the grandson of the Huang Di), is the progenitor of the Huang surname, which seems to conflict with the narrative that Da Lian is the ancestor of the Huang family. This perspective can be traced back to the famous scholar Wang Jian from the Southern Dynasties (420-589 CE) in his work *Xing Pu* (Genealogy of Surnames). However, this discrepancy may reflect differences in understanding across different historical periods and cultural contexts. To reconcile this conflict, scholars have proposed several hypotheses. One such hypothesis is that Lu Zhong may represent the matrilineal ancestor of the Ji surname during a matriarchal society, while the family of Bo Yi represents the patriarchal ancestors of the Ying surname after the transition to a patriarchal society (14).

The conclusions of this study seem consistent with this hypothesis. If Lu Zhong were the patriarchal ancestor of the Huang surname, then the Huang surname would not be genetically related to the Xu or Liang surnames, as no such records connecting Lu Zhong to Xu or Liang are found in the literature. However, the genetic relationship found between the Huang surname and the Xu or Liang surnames in our study here aligns with the Bo Yi family narrative as documented in *Shiji* (Figure 2). This leans towards supporting the conclusion that Da Lian is the patriarchal ancestor of the Huang surname. This further highlights the profound academic achievements demonstrated by Sima Qian in his monumental work *Shiji*.

There are numerous records about the Xia Dynasty in texts like *Shiji*, but due to skepticism from Western scholars and the temporary lack of archaeological evidence, there has yet to be a consensus in the academic community about whether the Xia Dynasty truly existed. A new, independent research perspective may provide a unique piece of evidence for validating the existence of the Xia Dynasty. According to *Shiji*, the Huang State and Xu State were established during the early Xia Dynasty as the fiefs of the two sons of Bo Yi, a prominent minister of the Xia Dynasty, and these states are said to be the origins of the Huang and Xu surnames. This study traced the genetic origins of these two surnames from the perspective of surname genetics, and the findings are largely consistent with the historical records of the Huang and Xu States. This provides strong genetic evidence supporting the existence of these two States and Xia Dynasty.

Although the overall sample size in our study is considerable, the sample size for individual surnames is relatively limited. As such, the conclusions of this study are expected to be further reinforced through future large-scale and more targeted systematic research. However, it is worth noting that the conclusions here have a high level of credibility because the three independent studies targeting different surnames all yielded consistent results that align with and mutually corroborate the historical records. If the smaller sample sizes had indeed caused any bias in the conclusions, such cross-surname consistency would be difficult to maintain. Furthermore, the majority of the sample data used in this study comes from 23Mofang, a company whose sample collection process did not specifically target any particular surname, thereby ensuring that the study’s conclusions are less likely to be influenced by bias.

In conclusion, our study here identified a genetic lineage of the Huang surname from Hubei that aligns with the historical records of the Huang surname’s origin. Additionally, genetic lineages related to surnames such as Xu and Liang were also identified, further supporting the historical records. The existence of these genetic lineages corroborates the historical accounts of the Huang and Xu States established during the early Xia Dynasty. Using surname genetics to independently verify the authenticity of the origins, transmission, and related dynasties, states, and historical figures may in the future produce many important results in molecular historical research.

## Acknowledgements

We thank Xiaozhou Liu and Zhiyi Xia for technical assistance. This work was supported by the National Natural Science Foundation of China (Grant No. 81171880) and the National Basic Research Program of China (Grant No. 2011CB51001).

## Author contributions

SH participated in the research design, data analysis, guidance of the team’s technical work, and manuscript writing.

## Ethical approval

All procedures performed in studies involving human participants were in accordance with the ethical standards of the institutional and/or national research committee and with the 1964 Helsinki declaration and its later amendments or comparable ethical standards.

## Supplementary Materials

**Supplementary Table S1. Haplotype distribution for the surnames Huang, Xu, and Liang.** Branches with fewer than 5 samples nationwide are not listed. For listed branches that have fewer than 5 samples nationwide in certain surnames, the data are not shown.

